# Proof of concept of a novel *ex vivo, in situ* method for MRI and histological brain assessment

**DOI:** 10.1101/2020.04.08.031682

**Authors:** Josefina Maranzano, Mahsa Dadar, Antony Bertrand-Grenier, Eve-Marie Frigon, Johanne Pellerin, Sophie Plante, Simon Duchesne, Christine L. Tardif, Denis Boire, Gilles Bronchti

## Abstract

MRI-histology correlation studies of the *ex vivo* brain mostly employ fresh, extracted (*ex situ*) specimens, aldehyde fixed by immersion. This method entails manipulation of the fresh brain during extraction, introducing several disadvantages: deformation of the specimen prior to MRI acquisition; introduction of air bubbles in the sulci, creating artifacts; and uneven or poor fixation of the deeper regions of the brain.

We propose a new paradigm to scan the *ex vivo* brain, exploiting a technique used by anatomists: fixation by whole body perfusion, which implies fixation of the brain *in situ*. This allows scanning the brain surrounded by fluids, meninges, and skull, thus preserving the structural relationships of the brain *in vivo* and avoiding the disadvantages of *ex situ* scanning. Our aims were: 1) to assess whether months of *in situ* fixation resulted in a loss of fluid around the brain; 2) to evaluate whether *in situ* fixation modified antigenicity for myelin and neuron specific marker; 3) to assess whether *in situ* fixation improved the register of *ex vivo* brain images to standard neuroanatomical templates in pseudo-Talairach space for morphometry studies.

Five head specimens fixed with a saturated sodium chloride solution (a non-standard fixative used in our anatomy laboratory for neurosurgical simulation) were employed. We acquired 3D T1-weighted (MPRAGE), 2D fluid-attenuated inversion recovery T2-weighted turbo spin echo (T2w-FLAIR), and 3D gradient-echo (3D-GRE) pulse sequences of all brains on a 1.5T MRI. After brain extraction, sections were processed for binding with myelin basic protein (MBP) and neuronal nuclei (NeuN) primary antibodies by immunofluorescence.

This study showed that all but one specimen retained fluids in the subarachnoid and ventricular spaces. The specimen that lost fluid was the oldest one, with the longest interval between the time of death and the MRI scanning day being 403 days. All T1-weighted images were successfully processed through a validated pipeline used with *in vivo* MRIs. The pipeline did not require any modification to run on the *ex vivo-in situ* scans. All scans were successfully registered to the brain template, more accurately than an *ex vivo-ex situ* scan and exhibited positive antigenicity for MBP and NeuN.

MRI and histology study of the *ex vivo-in situ* brain fixed by perfusion is feasible and allows for *in situ* MRI imaging for of at least 10 months post-mortem prior to histology analyses. Fluids around and inside the brain specimens and antigenicity for myelin and neurons were all well preserved.

## 1. INTRODUCTION

Neuroscientists studying the structure of the brain need histology to biologically validate their imaging findings, e.g. from magnetic resonance (MRI) ^1^. Physio- and anatomopathological insights into neurological diseases also require histological assessment alongside imaging of the *ex vivo* specimens, to draw conclusive interpretations regarding the nature of the underlying biological changes that are observed in the *in vivo* findings ^2-5^. For example, myelin content modulates the T1 relaxation times in MRI, which may be directly validated by immunohistochemical analysis of myelin basic protein (MBP) in the same scanned specimen ^6^.

Correlating MRI and histology of the same brain specimen is only possible *ex vivo* (post-mortem). The reliability of this correlation is contingent on two key points: 1) the distortion must be minimal between MRI and histology sections, to ensure that the spatial correspondence (registration) between modalities is maintained ^7^; and 2) brain tissue must be appropriately preserved, to ensure that specific antigenicity properties are retained, ^1^ i.e. the fixative solutions used do not impact or destroy the antigens that need to be measured.

Currently, most of the *ex vivo* brain MRI-histology studies employ fresh autopsy-extracted specimens (*ex situ*), aldehyde-fixed by immersion or frozen and kept in brain banks ^8^. However, these fixation approaches, which entail manipulation of the fresh brain during extraction, have major and inevitable disadvantages for the later MRI and histology stages of analysis, such as: 1) deformation of the freshly extracted tissue, which compromises the registration quality between histological and MRI images ^7^; 2) a gradient of fixation from the surface to the core of the organ caused by slow diffusion and/or uneven penetration of the fixative solution ^9^; 3) a need for special containers filled with a proton-free susceptibility matching fluid to acquire the MRI scans ^10^; 4) artifacts produced by air bubbles trapped in the subarachnoid spaces around the brain^10^; and 5) degradation of the superficial layers of tissue exposed to air ^10^.

An alternative method to *ex vivo-ex situ* imaging is *ex vivo-in situ* MRI scanning, i.e. with the brain remaining inside the head. This provides images in a similar setting to the *in vivo* case: a brain surrounded by its fluid, meninges, and skull, without deformation. In fact, fluids preservation in the subarachnoid and intraventricular compartments and the lack of deformation are hallmarks of *in situ* brain imaging ^2, 11^. To date, these studies are conducted only in unfixed post-mortem specimens in hospitals where MRI scanners are readily available within a few hours of the time of death ^2, 11^. Prior to histology or any subsequent treatments, the brain needs to be removed *en bloc*, thus manipulated in the fresh-unfixed state, introducing deformation to the organ, (*en bloc* = removal of both hemispheres, cerebellum and brainstem together after sectioning the cranial nerves, arteries, and proximal spinal cord, immediately distal to the foramen magnum) ^2, 11^. Consequently, deformation is prevented in the MRI scans, but not in the histology images that have to be registered to the MRI, not fully addressing the issue.

With respect to fixation methods for brain-banking, the two most prevalent methods are: cryopreservation at −80°C, preferred for studying brain biochemistry, and aldehyde-based chemical preservation, preferred for studying cellular characteristics ^8,12^. Formaldehyde perfusion through the vascular system is preferable to immersion, since the vessels deliver the fixative to deep organ areas ^13^, whereas immersion fixes the most superficial brain structures well, but the deepest areas may experience degradation, especially in larger specimens ^8 13-16^. Perfusion fixation is not common in autopsy procedures, which, as mentioned above, routinely remove the brain *en bloc* prior to fixation by immersion ^8^. Conversely, *in situ* perfusion techniques are the standard procedure of anatomical laboratories.

In the last thirty years, few studies have been performed using *in situ* brain fixation for histological study purposes ^17-19^. The main objective of these studies has been neurosurgical training, ^17^ plastination for anatomical teaching, and macroscopic gross anatomy dissection ^18, 19^. However, an *in situ* perfusion fixation technique provides equivalent or superior histology results compared to fixation by immersion ^8, 20-24^. One must consider however the potential dehydration with time, which could entail fluids loss around the brain, impacting the quality of an MRI scan.

The current study proposes a new paradigm to scan the *ex vivo* brain, exploiting the technique used by anatomists: perfusion fixation of the body, which implies *in situ* fixation of the brain ^17^. This allows scanning of the full head in a similar setting than the *in vivo* brain, avoiding manipulation and deformation of a freshly extracted brain at any point of the process, the need for a container for MRI, ^10^ and the occurrence of air artifacts and tissue degradation ^10^. Additionally, the perfusion method decreases the inevitable gradient of fixation produced by the immersion technique, providing more homogeneous images from the surface to the core of the organ ^9^. Finally, an *in situ*-perfused-fixed brain could be potentially conserved for many months before extraction of the organ for histological analysis, increasing the current *in situ* MRI window, which is only of a few hours post-mortem ^11^.

For this paradigm to stand, we set out to prove three fundamental concepts:

1. that the fluid around (subarachnoid compartment) and inside (intraventricular compartment) the brain was preserved after months of conservation;
2. that the antigenicity of the brain tissue was maintained even after months of *in situ* fixation using a non-standard brain fixative, such as a saturated sodium chloride solution; and
3. that our specimens *in situ-ex vivo* images would be effectively registered to standard neuroanatomical templates in pseudo Talairach space ^25^, for future morphometry studies.

To our knowledge, this study presents the first MRI and histology results of a sample of *ex vivo*-*in situ* perfused-fixed brains using a saturated sodium chloride solution as fixative.

## 2. METHODS

### 2.1. Population

We employed a convenience sample of five head specimens fixed by perfusion at the Anatomy Laboratory of the University of Québec in Trois-Rivières of donors not affected by neurologic or psychiatric diseases. The study was approved by the University’s Ethics Subcommittee on Anatomical Teaching and Research.

All head specimens were prepared (severed) prior to MRI scanning following the preparation guidelines used for neurosurgical simulation, which are: laminectomy of the cervical vertebrae C6 and C7, closure of the dural sac around the spinal cord using non-distensible cotton thread, transection of the spinal cord distal to the closed dural sac, followed by transection of the head at that same level (Figure 1 presents all steps involved in this new approach for *ex vivo-in situ* MRI and histology analysis). Head specimens were kept wrapped in a cotton sheet humidified with a standard conservation solution (water-based solution of glycerol and phenol).

**Figure 1:**
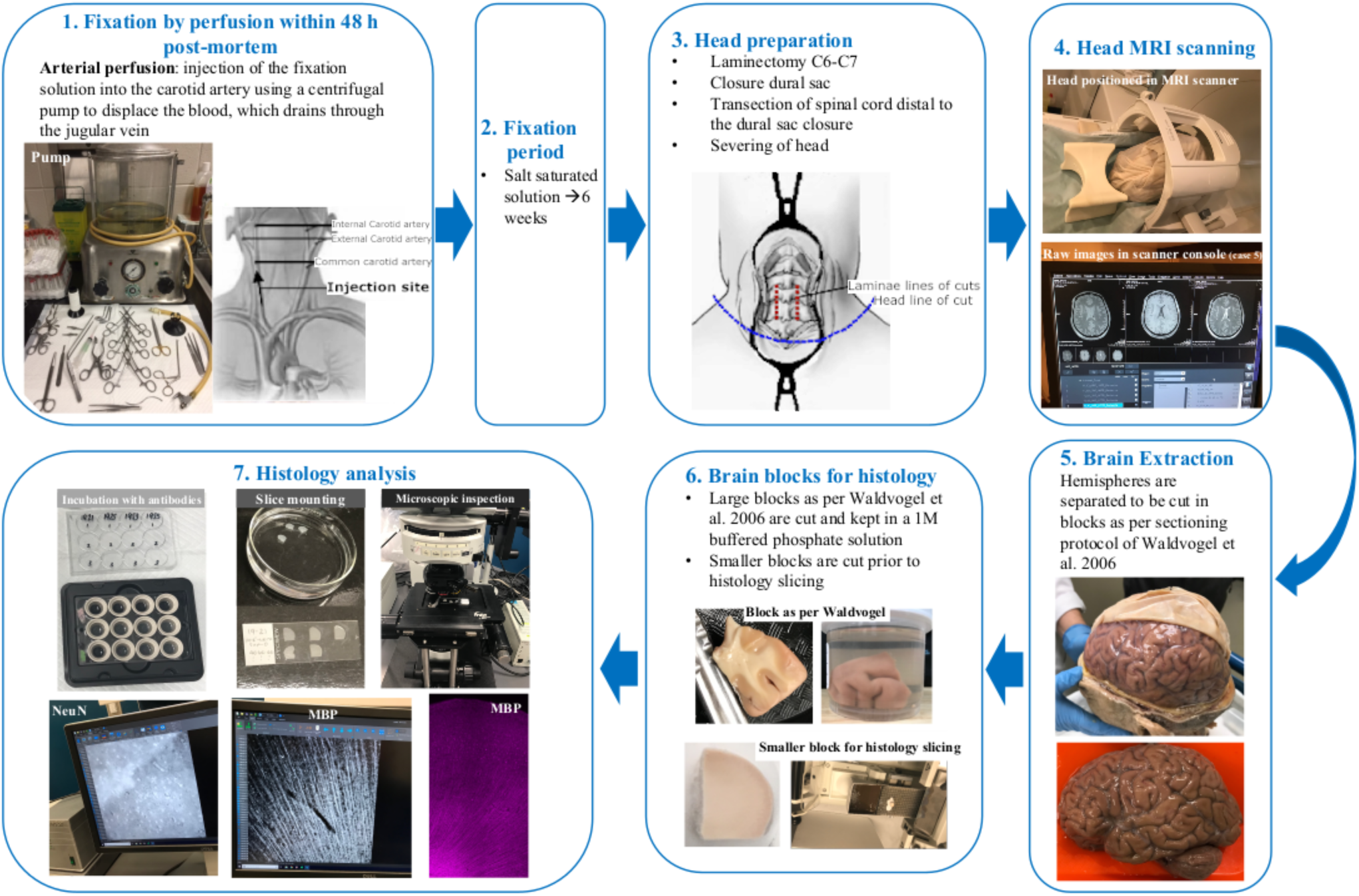
MRI-histology flow-chart

Table 1 shows demographic data, along with the intervals to fixation, head specimens’ preparation for scanning, brain extractions, and histology analysis.

**Table 1:** Demographic characteristics of the donors

| ID | Age at time of death | Sex | Time interval |  |  |  |  |  |
| --- | --- | --- | --- | --- | --- | --- | --- | --- |
|  |  |  | To fixation | To head preparation | To MRI scanning | Between preparation and MRI | To brain extraction | To histology analysis |
| 1 | 94 | F | 8 h | 231 d | 403 d | 172 d | NA | NA |
| 2 | 87 | F | 8 h 30 m | 126 d | 298 d | 172 d | NA | NA |
| 3 | 76 | M | 6 h | 166 d | 168 d | 2 d | 236 d | 242 d |
| 4 | 64 | M | 7 h 20 m | 145 d | 147 d | 2 d | 157 d | 221 d |
| 5 | 87 | M | 7.5 | 78 d | 81 d | 3 d | 83 d | 89 d |

F=female; M=male; h=hours; d=days; m=minutes; NA=not applicable.

### 2.2. Fixative solution

*Sodium chloride solution:* Each corpse was embalmed with 25 l of an aqueous saturated salt solution (36% NaCl) containing 0.8% formaldehyde (Fisher Scientific, 37% solution stabilized with 15% methanol); 0.72% phenol (Anachemia, 90% solution); 2% Glycerol (Fischer scientific); 16% isopropyl alcohol (AdC, 99%) ^26^. This solution produces less tissue retraction than the standard formaldehyde fixative solution at 4%. Therefore, the macroscopic aspect of brains at extraction is closer, in size and consistency, to that of a fresh brain. Additionally, previous studies using the same fixative have shown good cellular morphology in histology sections of the nervous tissue ^27^. To our knowledge, specific antigens have not been assessed in nervous tissue fixed in this way.

### 2.3. MRI acquisition

Two different MRI protocols were tested in as many sessions using a 1.5 tesla (T) Siemens MAGNETOM Avanto MRI scanner (Drummondville Hospital, Québec, Canada).

*Protocol 1*: Head specimens 1, 2, 3, 4 were scanned using the following pulse-sequences:

- 3D T1-weighted magnetization prepared rapid gradient echo sequence (MPRAGE).
- 2D fluid-attenuated inversion recovery sequence T2-weighted turbo spin echo sequence (T2w-FLAIR) ^28^.

*Protocol 2*: Specimens 2, 3, and 5 were scanned using the following sequences:

- 3D gradient echo (GRE) sequence with various flip angles, to allow the calculation of T1 relaxation times ^29^.
- 3D-GRE pulse-sequence with different echo times, to allow the calculation of T2* relaxation times ^29^.

The first protocol was used to assess and compare the performance of conventional MRI protocols used in clinical research (designed for *in vivo* scanning) ^2, 3, 30^ in the *ex vivo* setting. Figure 2 shows examples of *ex vivo* and *in vivo* T2w images acquired in a similar 1.5T MRI scanner, side by side. The second protocol aimed to assess the *ex vivo* performance of pulse sequences that allow quantitative evaluation, through the computation of T1 and T2* relaxation curves ^29^. Additionally, these 3D-GRE sequences allow the generation of synthetic T1w and synthetic T2*w images, which could potentially replace the need to acquire MPRAGE and standard T2w-FLAIR sequences ^29^ (Figure 3 presents the MPRAGE and synthetic T1w of one specimen).

**Figure 2:**
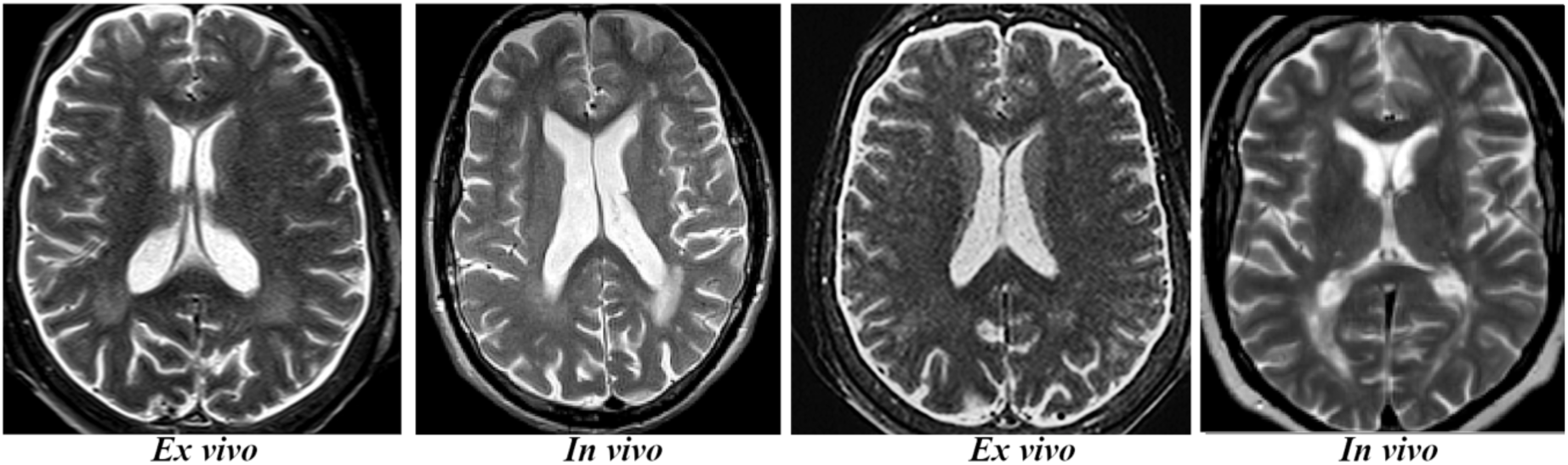
Examples of T2w-FLAIR cases *ex vivo* and *in vivo* showing similar morphological characteristics and fluid distribution in the subarachnoid and intraventricular compartments. In vivo images are of a living subject part of the *Alzheimer’s disease neuroimaging initiative (http://adni.loni.usc.edu)*.

**Figure 3:**
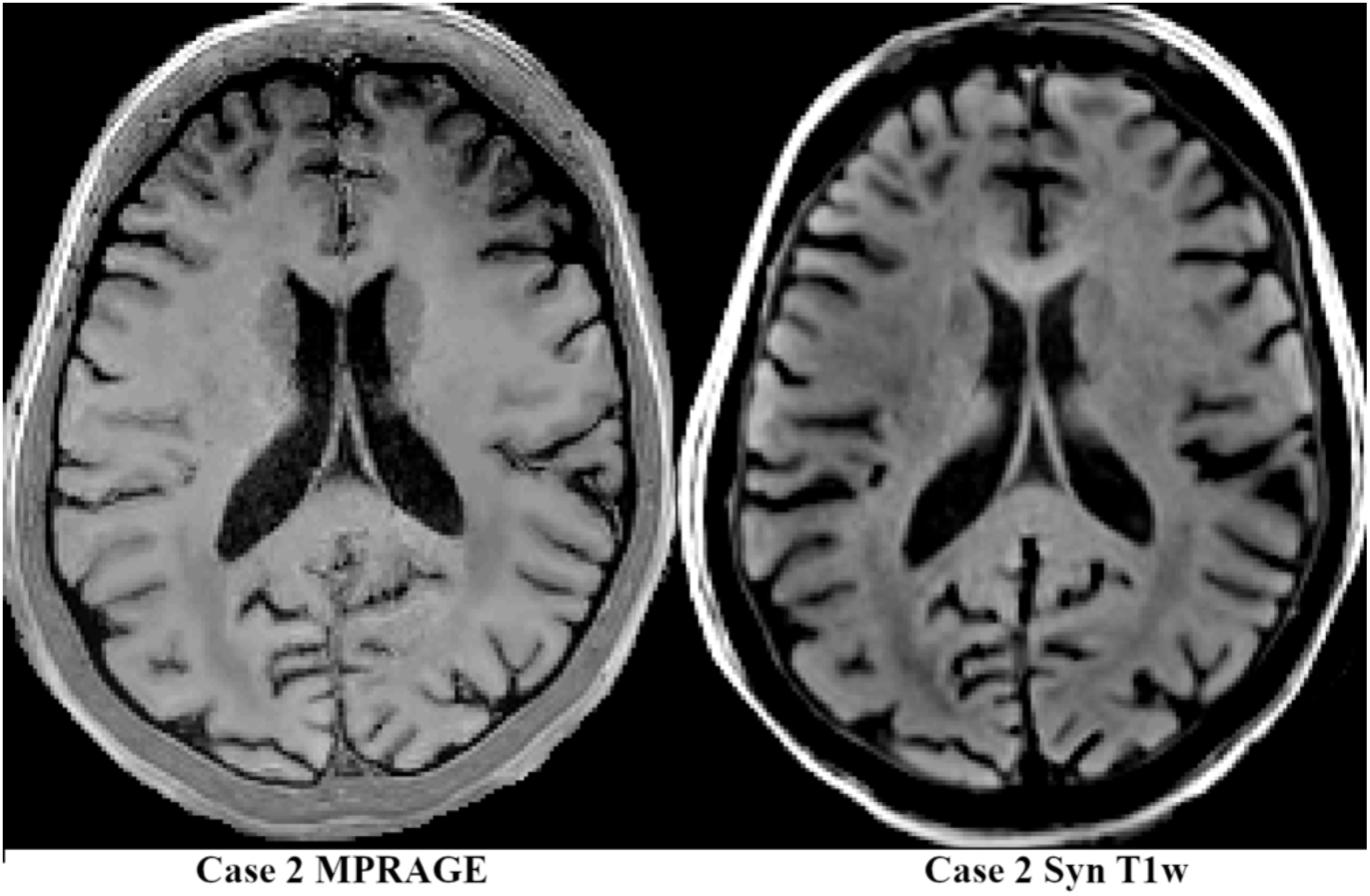
MPRAGE and synthetic T1w images generated using the 3D-GRE pulse sequences

The acquisition parameters of each pulse-sequence are listed in Table 2.

**Table 2:** Parameters of the pulse-sequences

|  | **3D-T1w**<br>**(MPRAGE)** | **2D T2w-**<br>**FLAIR** | **3D- GRE (1)** | **3D-GRE (2)** | **3D-GRE (3)** |
| --- | --- | --- | --- | --- | --- |
| **TR (ms)** | 2000 | 15000 | 45 | 45 | 45 |
| **TE (ms)** | 2.94 | 81 | 8 | 8 | 37 |
| **TI (ms)** | 900 | 3180 | NA | NA | NA |
| **FA (degrees)** | 9 | 180 | 7 | 35 | 7 |
| **BW (Hz/pixel)** | 180 | 110 | 210 | 210 | 210 |
| **GRAPPA**<br>**acceleration**<br>**factor** | NA | NA | 2 | 2 | 2 |
| **Voxel size (mm x**<br>**mm x mm)** | 1x1x1 | 1x1x1.5 | 1x1x1 | 1x1x1 | 1x1x1 |

ms=millisecond; mm=millimeter; NA=not applicable.

### 2.4. MRI image processing

Images were converted to MINC format using *dcm2mnc* tool, part of the Brain Imaging Centre minc-toolkit (http://www.bic.mni.mcgill.ca/ServicesSoftware/ServicesSoftwareMincToolKit). T1w and T2w images were preprocessed in three steps:

1. Denoising using *mincnlm* tool, also part of the minc-toolkit ^31^.
2. Image intensity nonuniformity correction using N3 (*nu_correct* tool, part of the minc-toolkit) ^32^.
3. Intensity normalization into range 0-100 using *volume_pol* tool (part of the minc-toolkit).

The T1w images were then both linearly (9 parameter registration, using bestlinreg_s2 tool),^33^ and nonlinearly (using *minctracc* tool), registered to the International Consortium for Brain Mapping of the Montreal Neurological Institute (MNI-ICBM)152 average brain template ^25, 34^. A brain mask was generated for the T1w image using *BeAST* ^35^.

Quality control images were generated for the linear and nonlinear registration steps by overlaying the contours of the MNI-ICBM152 template (depicted in red) on the T1w images after applying the obtained linear and nonlinear transformations, respectively (See section 2.6. and Figure S1 in the Supplemental Material document for details of the quality control images).

Synthetic T1w images were generated by linear subtraction of the higher and lower flip angle images (i.e. TE8 FA35 and TE8 FA7), based on the method proposed by Chen et al. 2018 ^29^. Both images were denoised prior to this step, and a λ value of 1.4 was used as the weight for the proton density weighted image.

The same image processing steps were performed for a T1w MRI of a conventional (fresh) extracted brain from the Allen Institute: MRI *ex vivo-ex situ* ^36^. (case H0351.1009, available at http://human.brain-map.org/mri_viewers/data)

### 2.5. Histological analysis

After the MRI sessions, the brains of specimens 3, 4 and 5 were extracted *en bloc*. Specimens 1 and 2 were not considered for histologic analysis due to: 1) loss of fluid surrounding the brain of specimen 1 (see Figure 4), and 2) insertion of latex in the arteries of specimen 2 (this specimen was intended for anatomy teaching of the vascular system). Subsequently, each hemisphere was sectioned in small blocks using a number 23 scalpel blade, as per the dissection protocol of Waldvogel et al (2006) ^1^, generating multiple approximately 3×3×3 cm^3^ blocks from the post-central, pre-central, superior temporal gyri and the occipital lobes. All blocks were kept in a 0.1M phosphate buffered solution (pH 7.3) prior to histological analysis.

**Figure 4:**
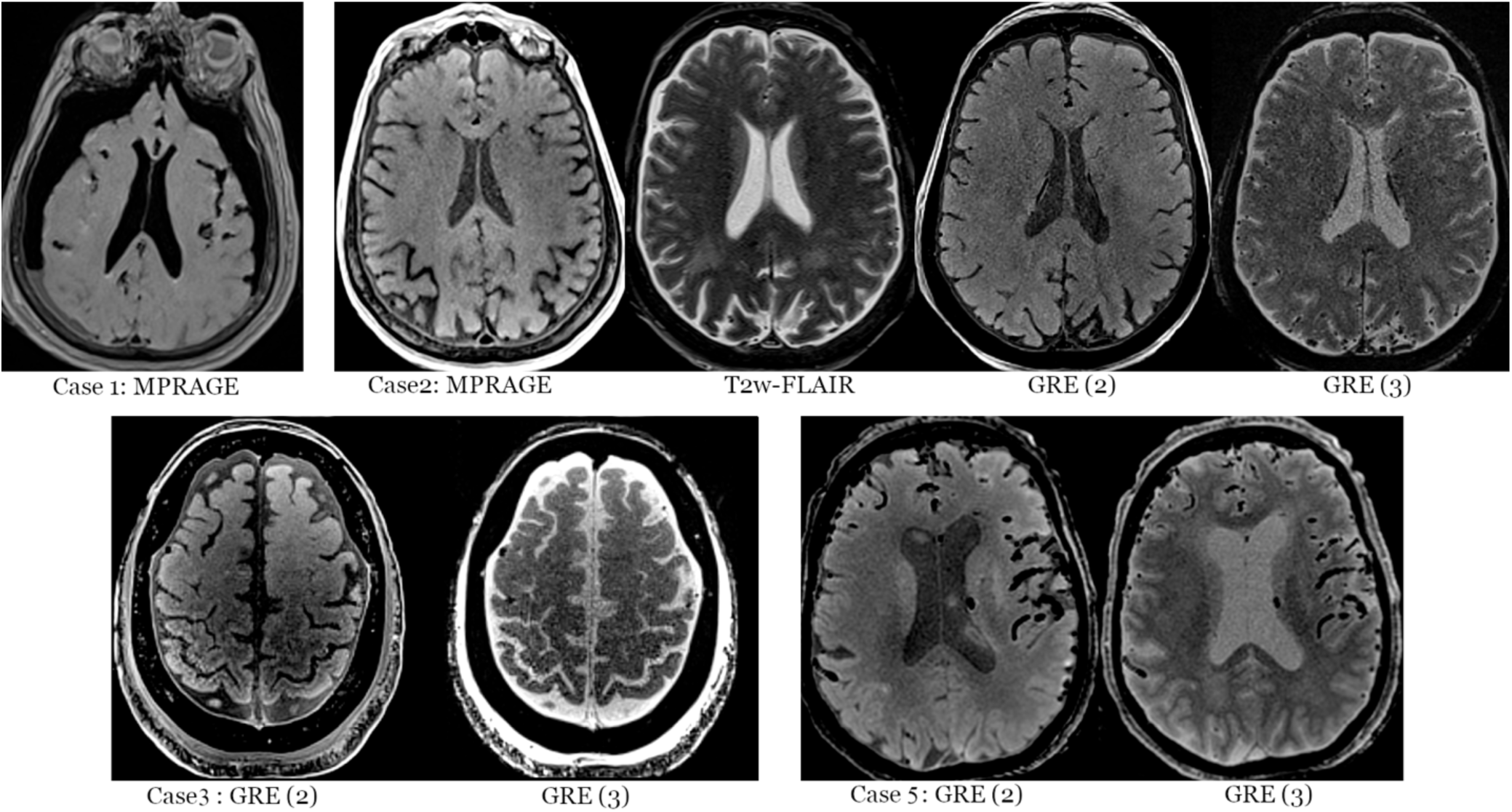
MRI examples of different cases. Note that Case 1 shows lack of fluids in the subarachnoid and ventricular spaces.

At the time of histological analysis, a random block of each specimen was used to assess two antigens: myelin basic protein (MBP) and neuronal nuclei (NeuN), by immunofluorescence. A smaller piece of approximately 1×1×1 cm^3^ was cut from each randomly selected block and placed it in a 30% 0.1M phosphate buffered sucrose solution until sinking (48 to 72h). Then, the blocks were frozen with dry-ice and kept at −80°C until cutting them in 40- and 60-µm sections using a microtome. Subsequently, slices were rinsed in a 0.1M phosphate buffered solution, a 50% methanol solution, and again in a 0.1M phosphate buffered solution, then soaked overnight in blocking solution (1% Normal donkey serum, 2% Triton, 0,1M Glycine). Finally, all slices were incubated with MBP (1:1000 rabbit anti-MBP, Abcam; ab40390) and NeuN (1:6000 guinea pig anti-NeuN, Millipore Sigma, ABN90) primary antibodies (72 h incubation at 37°C), then rinsed using 0.1M phosphate buffered solution, and incubated with the corresponding secondary antibodies: 1:500 anti-rabbit alexa 555 (Biolegend;406412) and 1:500 anti-guinea pig alexa 647 (Jackson Laboratories;706-605-148).

All the slices were visualized and photographed using a DSU Spinning Disk confocal microscope (Olympus BX51W1) coupled to the Neurolucida software (MBF Bioscience).

### 2.6. Statistical, MRI, and histological analyses

Demographic variables and post-mortem intervals to fixation, head specimen preparation, scanning, brain extraction and histologic analyses are presented using descriptive statistics, as either mean and standard deviation (SD), or median and range, according to their distribution.

The MRI registrations to the ICBM-MNI model were assessed qualitatively through visual quality control using a viewing panel with 60 images extracted from axial, sagittal, and coronal slices (20 each) throughout the resampled brain volume, with the contours of the ICBM152 template overlaying each slice ^33^ (See Figure S1 in the Supplemental material document).

The histology analysis for presence or absence of myelin and neuronal antigenicity was performed quantitatively, in a binary fashion, as either positive or negative antigenicity as depicted by immunofluorescence.

All results were computed using SPSS version 24.0 (IBM Corporation, Armonk, New York).

## 3. RESULTS

The mean age at the time of death of the donors was 81.6 years old, with a standard deviation (SD) of 11.8. The male to female ratio was 3:2. The causes of death of the donors were: sudden death by cardiac arrest, cardiac insufficiency, pneumonia, larynx carcinoma, and lung carcinoma. None of the donors had a reported neurodegenerative, neurological, neurodevelopmental, or psychiatric disease.

The mean interval in hours (h) between the time of death and fixation was 7.5 h (SD: 0.9). The mean interval in days (d) between the time of death and head preparation was 149.2 d (SD: 56.1); the mean interval between the time of death and MRI scanning was 219.4 d (SD 129.3). The median interval between the time of head preparation and the MRI scanning session was 3 d (range: 2-172); the mean interval between the time of death and the time of brain extraction was 158.7 d (SD: 76.5); and the mean interval between the time of death and the time of histological analysis was 184 d (SD: 82.9).

Out of the five scanned specimens, only one showed poor fluid preservation around the brain and in the ventricular system, namely, specimen number 1, which was the oldest conserved specimen, with the longest time interval between the time of death and the MRI scanning day being 403 d, and the longest interval between the time of death and head preparation, 231 d (See Figure 4).

Specimens 2 to 5 showed a good preservation of fluid around the brain and in the ventricular system, comparable to the distribution of fluid in an *in vivo* setting (See Figures 2, 3 and 4). All the specimens histologically analysed (specimens 3 to 5) exhibited positive antigenicity for MBP and NeuN (See Figure 5).

**Figure 5:**
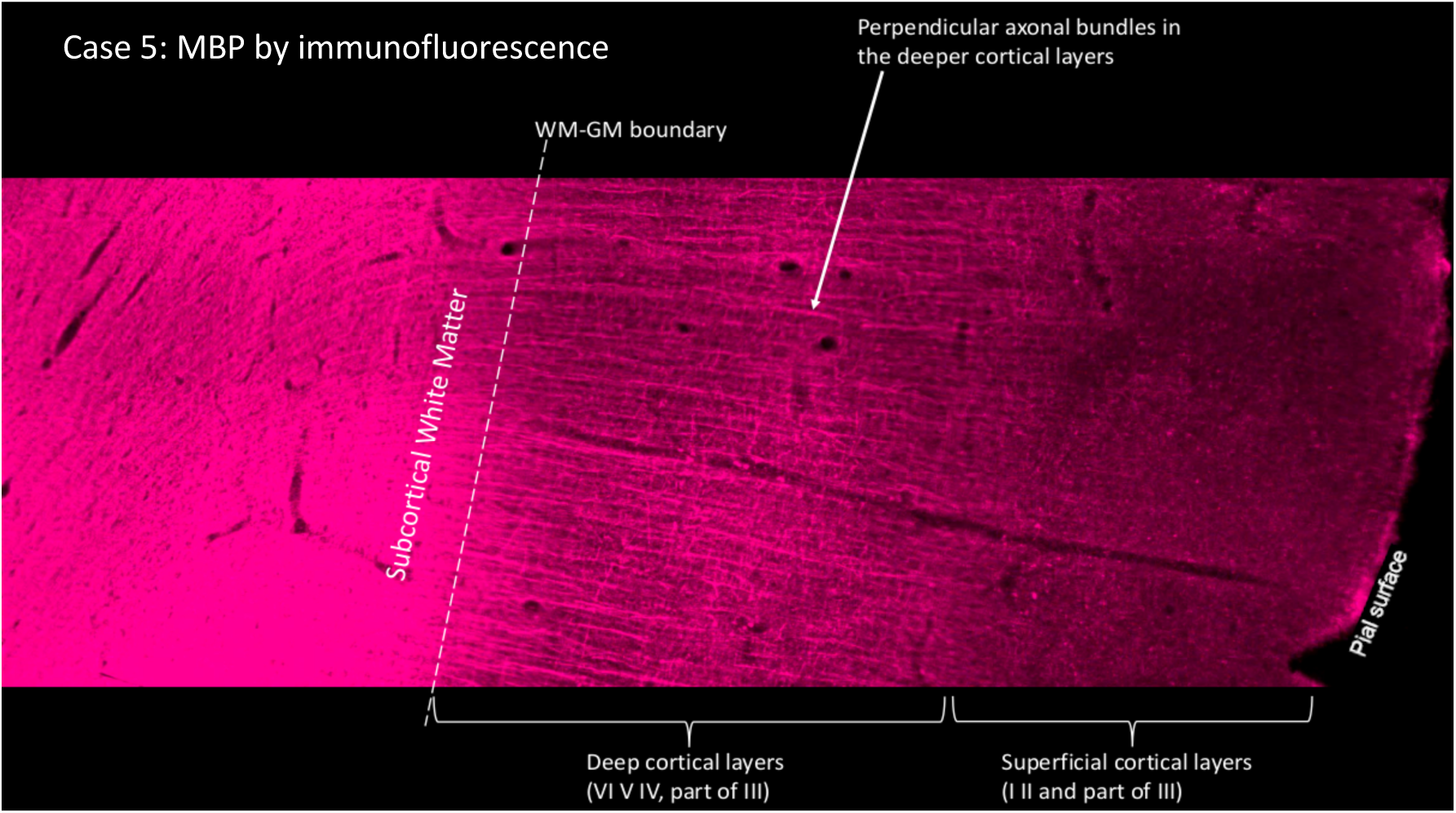
Example of Immunofluorescence staining for myelin basic protein (MBP). Note the change of color density (myelin density) decreasing from the subcortical white matter towards the pial surface.

The T1w images of all acquired MRI scans (bar number 1, due to fluid loss) were successfully processed using an already validated pipeline applied in *in vivo* MRI scans ^33^. The pipeline did not require any modification to successfully run on the *ex vivo* scans.

All scans were successfully registered to the MNI ICBM 152 brain model, more accurately than the Allen Institute *ex vivo-ex situ* MRI scan, which exhibited loss of normal ventricular volume, creating a geometrical distortion that precludes a good fitting to the average brain model (see in Figure 6 how the red contour cannot follow the pial, nor the ventricular surface of the *ex vivo-ex situ* scan).

**Figure 6:**
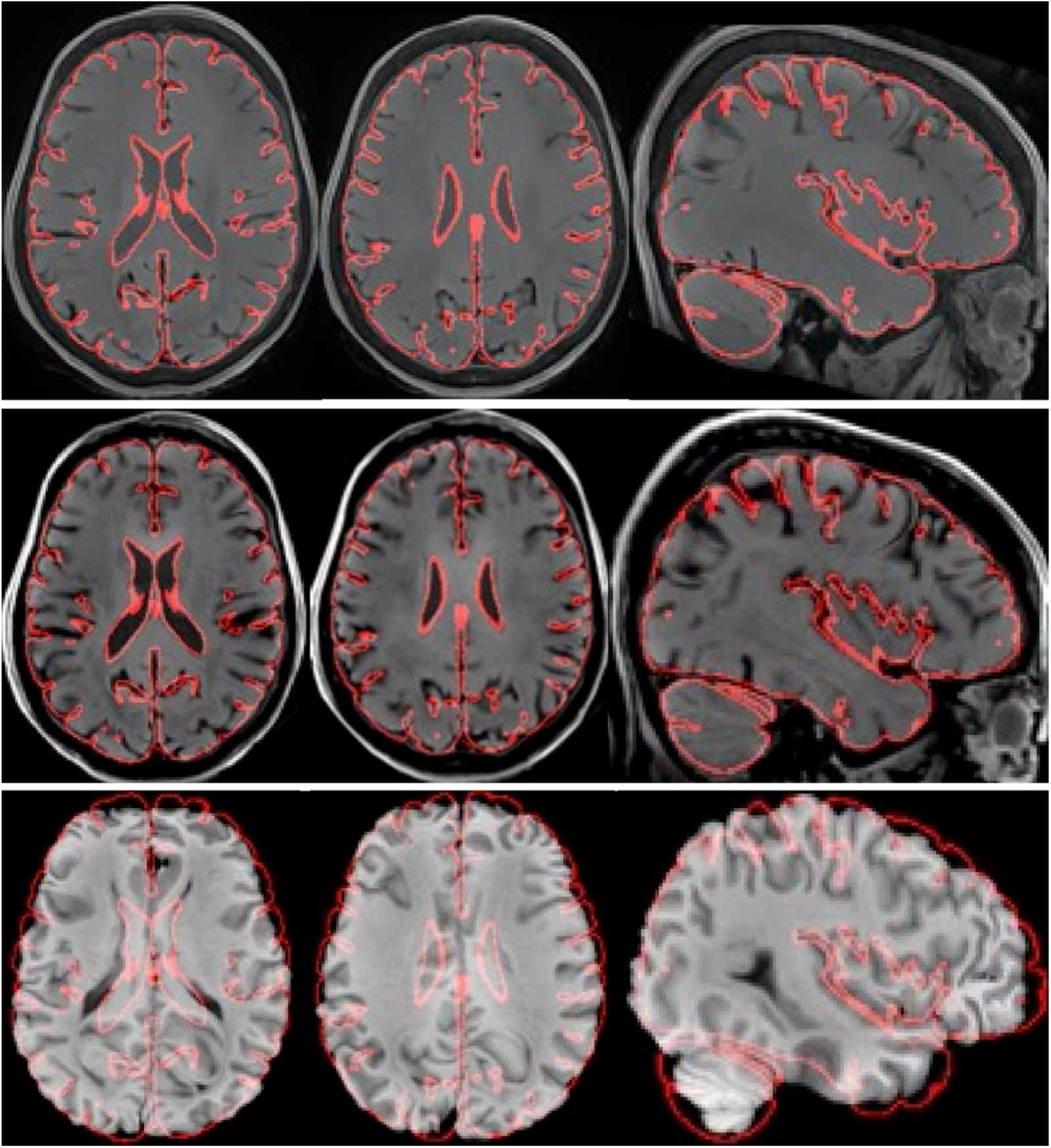
Comparison of non-linear registration to the ICBM MNI brain model of one of our *ex vivo-in situ* specimens (top panel: Syn T1w and middle panels: MPRAGE) vs. the non-linear registration to the ICBM MNI brain model of an *ex vivo-ex situ* specimen from the Allen Institute (bottom panel). Note how the red contour (representing the ICBM MNI brain edges) follows closely the edges of the *ex vivo-in situ* specimen, but not the edges of the *ex vivo-ex situ* brain.

## 4. DISCUSSION

The study of the *ex vivo* brain through histology and MRI is of paramount importance to accurately interpret *in vivo* MRI findings ^3-5^. For example, bright areas are present in the white matter of individuals with mild cognitive decline, but the nature of those changes cannot be ascertained, unless an histological analysis is performed, determining whether axons are injured, or myelin has been compromised, in comparison to the white matter of normal controls or the same individual in a different undamaged area ^3^. By establishing these correlations in *ex vivo* specimens, we can then infer the meaning of those MRI bright areas when we encounter them in the scan of a living individual.

In order to perform these types of MRI-histology correlation studies with enough statistical power, numerous *ex vivo* brain specimens affected by different pathologies, as well as matched normal controls, need to be available. This has led to the creation of brain banks around the world ^8^, which preserve brain samples according to different protocols: either as full hemispheres fixed by immersion in a formaldehyde solution, or slices of brain hemispheres cryopreserved at −80°C without fixation. Some brain banks have also explored the fixation by perfusion through the internal carotid and vertebral arteries ^8^. However, they perform the perfusion after the extraction of the fresh brain along with the vessels, having the advantage of better fixation of the deep regions, but not avoiding the manipulation and deformation of the freshly extracted brain. Additionally, the number of brains kept in banks may not suffice for the increasing needs of neuroscientists, who would then be forced to adapt their experiments to the number of samples available at the time. Moreover, the process of donation of the brains is also dependent on the possibility of performing an autopsy extraction within the first 48 hours post-mortem, which is frequently not available ^11^. Our novel approach to study the *ex vivo-in situ* brain proposes to solve some of the disadvantages of existing methods (e.g. deformation due to manipulation of the unfixed brain) and, at the same time, it has the potential of becoming a new source to obtain brain specimens for neuroscientific research projects. Our project takes advantage of the fixation method used by anatomists ^15, 17, 27^, which by perfusion of the whole body allows the fixation of the brain *in situ*. Our anatomy lab performs all the perfusions within the first 48 hours post-mortem, ensuring a minimal decay of tissues prior to fixation. Perfusion-fixed bodies can be used for many months for teaching purposes, including surgical and neurosurgical simulations ^17^, without macroscopic decay of the tissue qualities. Our anatomy lab employs different fixative solutions to fulfill specific aims (mainly saturated salt fixation for neurosurgical simulation purposes), so our pilot study aimed to test this solution that, based on our prior teaching experience, preserves the brain and meninges well in terms of volume and consistency. The salt saturated solution ^26^ is not standardly used in most anatomy labs, but has been shown to preserve cell morphology of various tissues well, including nervous tissue ^26, 27^.

Our *in situ* MRI protocol allowed assessment of the brain in a setting similar to the *in vivo* compartments of the head: brain surrounded by meninges and fluid in the subarachnoideal and intraventricular compartments (Figures 2, 3 and 4), which is essential to ensure lack of geometrical deformation of the organ. Only one of our specimens, the oldest specimen (more than one year of preservation) failed to retain the fluids intracranially, but all the other specimens did. The *in situ* situation allowed the use of imaging processing tools currently in place for *in vivo* scans ^33^, without the need of any software adaptation or changes. The quality and accuracy of post-processed *ex vivo-in situ* scans (e.g. scans registered to the standard space of the ICBM MNI brain model) was superior to the results obtained using *ex vivo-ex situ* scans ^36^, which experience deformation (see Figure 6 showing an *ex vivo-ex situ* case with collapsed ventricles).

Finally, there are potential additional advantages in using our *ex vivo-in situ* approach:

1. if a donor has pre-mortem MRI scans available, those scans could be potentially treated with similar imaging processing tools, allowing a better translation/registration of *in vivo-*to*-ex vivo* findings;
2. the preservation of a fixed brain *in situ* allows for many months of imaging using different MRI scanners operating at different field strengths, prior to extraction of the brain for histological analyses, allowing an easier translation of findings from 1.5T-to-3T-to-7T MRI analyses techniques;
3. the extraction of the brain is performed after fixation, so the organ is never manipulated fresh, which is when the organ is more friable. This would also allow (if need be) imaging the brain *ex situ*, avoiding the geometrical changes (e.g. collapse of the ventricles), and cutting the brain in blocks for subsequent histology, in a more precise fashion, since the tissue is not friable. Regarding histological analyses, all of our samples showed positive antigenicity using immunofluorescence for the two tested antigens: MBP and NeuN, proving that a saturated salt solution does not alter or destroy these antigens.

Since our study was a proof-of-concept investigation, it is not without limitations, all of which need to be addressed in future experiments:

- exploration of the preservation of the fluids in specimens between 298 and 403 days of interval between the time of death and the MRI scanning, since that seems to be the interval of time in which fluid preservation is compromised;
- acquiring MRI scans of the tissue blocks obtained following the protocol of Waldvogel et al. ^1^ is needed to allow a precise registration of the histology slice to the MRI of the block, and then the registration of the MRI of the block to the full brain MRI. These steps will allow a precise registration of histology slices to the full brain MRI.

Regarding the histology analyses, future work will address the following current limitations:

- testing other antigens, such as glial fibrillary acidic protein to assess astrocytes, CD11b, CD45, Iba1 to assess microglia ^37^, and also elements like iron,^38^ that tend to accumulate in the aging brain and in many neurodegenerative processes, using Perl Prussian blue ^38^; and
- performing analysis using immunohistochemistry ^37^, as opposed to immunofluorescence, to allow the preservation of the histology material for future revisions of the work, and a more precise quantification (e.g. semi-automatic color-based extraction method, which provides results expressed as chromogen positive pixels/mm^2^) ^37^.

## Conclusion

An MRI and histology study of the *ex vivo-in situ* perfusion-fixed brain is feasible and allows for *in situ* MRI imaging during at least 10 months post-mortem prior to histology analyses. Fluids around and inside the brain specimens are well preserved, keeping the geometry of the brain undistorted, and antigenicity for myelin and neurons is present.

The proposed method has the potential of changing the way in which post-mortem MRI and histology studies are performed.

## Data availability statement

The MRI data of this study is available from the corresponding author upon request. The code that supports the results of this study are available at https://github.com/BIC-MNI/minc-toolkit-v2.

