## Supplemental Material for "Proof of concept of a novel *ex vivo, in situ* method for MRI and histological brain assessment"

**SUPPLEMENTAL MATERIAL: A novel *ex vivo-in situ* approach to study the brain through MRI and histology: results of a pilot study**

**Figure S1:** example of the 60 image-panel created to QC the registration of the case to the MNI ICBM model

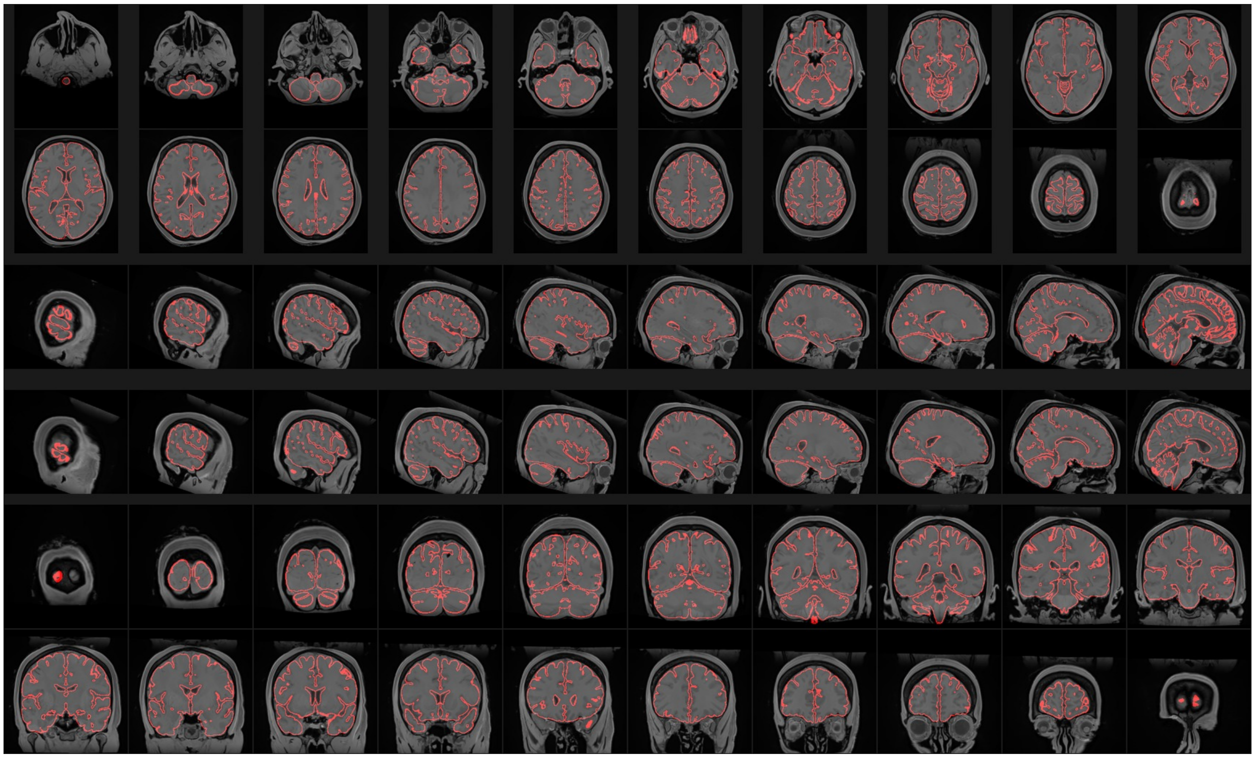
